# Altered connectome topology in newborns at risk for cognitive developmental delay: a cross-etiologic study

**DOI:** 10.1101/2024.06.20.599853

**Authors:** Anna Speckert, Kelly Payette, Walter Knirsch, Michael von Rhein, Patrice Grehten, Raimund Kottke, Cornelia Hagmann, Giancarlo Natalucci, Ueli Moehrlen, Luca Mazzone, Nicole Ochsenbein-Kölble, Beth Padden, SPINA BIFIDA STUDY GROUP ZURICH, Beatrice Latal, Andras Jakab

## Abstract

The human brain connectome is characterized by the duality of highly modular structure and efficient integration, supporting information processing. Newborns with congenital heart disease (CHD), prematurity, or spina bifida aperta (SBA) constitute a population at risk for altered brain development and developmental delay (DD). We hypothesize that, independent of etiology, alterations of connectomic organization reflect neural circuitry impairments in cognitive DD. Our study aim is to address this knowledge gap by using a multi-etiologic neonatal dataset to reveal potential commonalities and distinctions in the structural brain connectome and their associations with DD. We used diffusion tensor imaging (DTI) of 187 newborns (42 controls, 51 with CHD, 51 with prematurity, and 43 with SBA). Structural weighted connectomes were constructed using constrained spherical deconvolution based probabilistic tractography and the Edinburgh Neonatal Atlas. Assessment of brain network topology encompassed the analysis of global graph features, network-based statistics, and low-dimensional representation of global and local graph features. The Cognitive Composite Score of the Bayley Scales of Infant and Toddler Development 3^rd^ edition was used as outcome measure at corrected 2 years for the preterm born individuals and SBA patients, and at 1 year for the healthy controls and CHD.

We revealed differences in the connectomic structure of newborns across the four groups after visualizing the connectomes in a two-dimensional space defined by network integration and segregation. Further, ANCOVA analyses revealed differences in global efficiency (p < 0.0001), modularity (p < 0.0001), mean rich club coefficient (p = 0.017) and small-worldness (p = 0.016) between groups after adjustment for postmenstrual age at scan and gestational age at birth. Moreover, small-worldness was significantly associated with poorer cognitive outcome, specifically in the CHD cohort (r = -0.41, p = 0.005).

Our cross-etiologic study identified divergent structural brain connectome profiles linked to deviations from optimal network integration and segregation in newborns at risk for DD. Small-worldness emerges as a key feature, associating with early cognitive outcomes, especially within the CHD cohort, emphasizing small-worldness’ crucial role in shaping neurodevelopmental trajectories. Neonatal connectomic alterations associated with DD may serve as a marker identifying newborns at-risk for DD and provide early therapeutic interventions.

## 1 Introduction

The human brain connectome is characterized by the duality of highly modular structure and efficient integration, supporting specialization and efficient information processing (van den Heuvel & Sporns, 2019). Neuroimaging has used methods from graph theory and related fields to quantify these attributes of the brain connectome in health and disease (Fornito et al., lmno; Sporns, lmnn; Sporns et al., lmmo). More recently, a framework has been proposed that conceptually maps the human brain connectome along the two dimensions of segregation, which describes the modular structure and integration. The selection of these features as the two major dimensions stems from the observation that they capture fundamental aspects of the connectome’s organization, offering insights into the balance between local specialization (segregation) and global integration, also called small-worldness. Moreover, cognitively meaningful individual variation might be linked to how the brain tends to keep network costs down through segregated systems but at the same time promotes global, integrated processing (van den Heuvel & Sporns, 2019). In their framework, the morphospace defines a space where efficient organization occurs at the optimal trade-off between segregation and integration and depicts a biologically grounded, lower-dimensional representation of the complex brain connectome data originating from neuroimaging studies.

The duality of segregation and integration in the brain connectome emerges as a result of predominantly genetically programmed events during pre- and early postnatal development in humans. These attributes can be measured using MRI in fetuses, prematurely born newborns and young infants (Cao et al., 2017; Jakab, 2019; Ouyang et al., 2017). However, there is a scarcity of studies that specifically explored how this duality in brain neural circuitry is affected in the case of neurodevelopmental disorders. From the perspective of the morphospace framework, such neurodevelopmental (ND) disorders might introduce a shift of an individual’s connectome away from the optimal balance, placing it in a ‘suboptimal’ region. Among the many etiologies that may affect brain development during critical periods are neural tube defects, prematurity, and severe types of congenital heart disease (Bernier et al., 2010; Frey & Klebanoff, 2016; Little & Elwood, 2019; van der Bom et al., 2011). Children with congenital heart disease (CHD) (Latal, 2016), open spina bifida (spina bifida aperta, SBA) (Hepp et al., 2021) or those born prematurely (PB) (Wallois et al., 2020) face an increased risk of developing ND disorders. Among these, cognitive developmental delay (DD) is a clinical concern (Marino et al., 2012; Potijk et al., 2013; Ramsundhar & Donald, 2014), which is defined by deviations from meeting the expected cognitive developmental milestones (Petersen et al., 1998). The potential reason why these etiologies might develop DD is multifactorial and is likely a consequence of abnormal fetal cerebral blood flow (in CHD and SBA), low birth weight, young gestational age at birth, structural changes in SBA (Hee Chung et al., 2020; Hepp et al., 2021; Howell et al., 2019; Vonzun et al., 2023), and many more. While children with these etiologies may exhibit many similarities in cognitive developmental profiles, the extent to which they may share impairments in whole-brain network-level connectivity, and whether such impairments are associated with cognitive DD, remains incompletely understood.

Our knowledge in the field of connectomics mostly stems from case-control studies investigating a single pathology. Topological properties measured by graph-theory encompass both global characteristics, providing insights into the overall network structure, and local characteristics offering details about specific regions or nodes within the network. Key topological connectomic properties, such as integration, segregation, and rich-club coefficients – which refer to hubs (nodes with many connections) that are more densely interconnected with each other compared to other nodes – serve as reliable indicators for distinguishing between normal and abnormal brain networks (Lo et al., 2010; Owen et al., 2013; Shu et al., 2011). However, the existing neonatal literature shows inconsistent findings. In prematurity, for instance, studies by Zhao et al. (2019) and Kline et al. (2021) report decreased segregation compared to neonates with a higher gestational age at birth, while Sa de Almeida et al. (2021) observe an opposing trend with increased segregation in very preterm born infants (GA < 32 weeks) measured at term equivalent age. In terms of integration, a consistent decrease in prematurity compared neonates with a higher gestation age at birth is reported across studies by Zhao et al., (2019), Kline et al., (2021), and Sa de Almeida et al. (2021). Furthermore, the rich-club coefficient, indicative of highly interconnected regions, in the neonatal period is reported to be reduced in very preterm born infants (GA < 32 weeks) by Sa de Almeida et al. (2021) and increased in very preterm born infants (GA < 33 weeks) according to (Kim & Min, 2020). In CHD, studies reported heterogeneous results. Feldmann et al. (2020) and Schmithorst et al. (2018) find decreased integration during the neonatal period compared to healthy controls, while Ní Bhroin et al. (2020) observe no significant differences. Further, Feldmann et al. (2020) report higher segregation in CHD cases compared to healthy controls. As of the present, the spina bifida cohort remains unexplored by connectomic techniques, even though they could help to better understand network-level connectivity in this population.

Our hypothesis is that healthy developing neonatal structural connectomes occupy a more optimal region within the ‘integration-segregation morphospace’ compared to structural connectomes obtained from neonates at risk for cognitive DD. This would mean that newborns with specific etiologies and lower cognitive outcome scores within these etiologic groups might be associated with characteristic ‘disease zones’ in such representation of their connectomes. Therefore, we first aim to quantify alterations at the whole-brain network level, specifically analyzing integration, segregation, and rich-club coefficients in a single institutional, cross-etiologic cohort consisting of MRI data of CHD, SBA and PB infants compared to healthy controls and prove that these measures are associated with cognitive DD assessed at early childhood. We then aim to explore if local graph theory features might reveal additional insight into potentially identified disease zones by embedding the whole-brain connectomes’ local features in a lower-dimensional space.

## 2 Methods

### 2.1 Study Populations

The data used for this analysis originate from the following three cohort studies conducted at the University Children’s Hospital of Zurich: The “Heart and Brain” (HB) study (Meuwly et al., 2019), prospective collection of clinical cases with spina bifida undergoing fetal repair - “Spina Bifida” (SB) study (Moehrlen et al., 2020) and part of “The Swiss EPO Neuroprotection Trial” (EPO) (O’Gorman et al., 2015).

In the prospective HB study, neonates with CHD who required corrective or palliative cardiac surgery in the first weeks of life at the University Children’s Hospital Zurich from December 2009 to March 2019 were considered for inclusion (Feldmann et al., 2020; Meuwly et al., 2019). Infants were enlisted for participation in the study upon their admission to the University Children’s Hospital. Neonates were excluded if they had an identified or suspected genetic disorder. The infants who were part of the HB study underwent both pre- and post-surgical brain MRI. Additionally, a healthy control group was recruited between January 2011 and April 2019 from the post-natal ward at the University Hospital Zurich. For our present DTI study, only the pre-surgical MRI was analyzed, resulting in 67 neonates with CHD and 43 healthy controls. For more information about the study population see Meuwly et al. (2019).

In the EPO study, preterm born neonates aged between 26 and 31 gestational weeks were eligible for enrollment during the first 3 hours after birth (Natalucci et al., 2016; O’Gorman et al., 2015). In the randomized, double-blind placebo-controlled intervention study, Erythropoietin, a haematopoetic cytokine that is widely used for the treatment of anaemia in premature infants, was used as intervention for the PB infants. The exclusion criteria for the EPO study included the following: existence of a genetically defined syndrome, severe congenital malformation negatively impacting life expectancy, and severe congenital malformation adversely affecting prior palliative care or neurodevelopment (Fauchère et al., 2015). Structural and diffusion MRI (DTI) at term equivalent age was acquired. For the EPO study, only infants where the DTI scans were acquired in Zurich were included. For the present analysis of DTI data, a dataset of 58 neonates were included. More details about this study can be found in O’Gorman et al. (2015).

In the prospective SB study, neonates with open spina bifida who underwent open prenatal surgical repair between June 2014 and June 2020 at the Zurich Center for Fetal Diagnosis and Therapy were included. The MOMS inclusion and exclusion criteria were applied with some modifications to identify subjects eligible for fetal surgery (Moehrlen et al., 2023). Neonatal structural and diffusion MRI was acquired on a 1.5T and 3.0T clinical scanner. For this study, only data from the 3.0T scanner was utilized; to increase the homogeneity of the imaging data, subjects scanned at the 1.5T scanner were excluded. We only included cases if the DTI data quality was satisfactory, no brain lesions were present, and lateral ventricle size was 20 mm at maximum. This results in an SBA dataset of size 44.

In summary, for the present MRI study, subjects from these three studies were included (n_Controls_ = 43; n_CHD_ = 67; n_PB_= 58; n_SBA_ = 44) when good quality MR (T2-weighted and diffusion weighted) images existed (n_Controls_ = 42; n_CHD_ = 54; n_PB_ = 53; n_SBA_ = 44), and either a 1 or a 2-year developmental outcome was performed (n_Controls_ = 36; n_CHD_ = 52; n_PB_ = 52; n_SBA_ = 44). Further, subjects were excluded due to artifacts arising during image processing (n_Controls_ = 36; n_CHD_ = 49; n_PB_ = 50; n_SBA_ = 43), resulting in a total of 187 subjects from which 178 also have developmental outcome data (see supplementary Figure 1 and Figure 2 for a more detailed flow chart of the inclusion and exclusions; see supplementary Table 3 for the CHD types). All parents or caregivers gave written informed consent for the further use of their infants’ data in research. The ethical committee of the Canton of Zurich approved the studies for collecting and analyzing clinical data retrospectively (2016-01019, 2021-01101 and 2022-0115). The clinical variables in our work are based on a data registry that was established to collect all pertinent data in a prospective and systematic way. All the clinical descriptive data of the spina bifida newborns used in this study, were obtained from this REDCap™-based repository. The HB study was approved by the ethical committee of the Kanton Zurich, Switzerland (KEK StV-23/619/04). The study was carried out in accordance with the principles enunciated in the Declaration of Helsinki and the guidelines of Good Clinical Practice. The EPO study protocol was approved by the Swiss drug surveillance unit and the local ethical committees.

### 2.2 Magnetic resonance imaging (MRI) acquisition

The EPO cohort was scanned on a 3.0 T GE HD.xt MRI scanner (GE Medical Systems) using an 8-channel head coil. This system underwent a major upgrade replacing the gradient coils (GE Signa MR750). The upgraded scanner with the same head coil was used for the neonatal MRI scans from the HB study, including both CHD neonates and healthy controls, and the SB study. While the HB and EPO cohorts were scanned under natural sleep, the SBA infants were sedated using chloral hydrate.

The structural images used in our study were axial, coronal, and sagittal T2-weighted sequences. The structural, T2-weighted scanning protocol of the HB cohort included a fast-spin-echo sequence in axial, coronal and sagittal planes (FSE, voxel dimensions = 0.7 x 0.7 x 2.7 mm, repetition time (TR) = 3910 ms, echo time (TE) = 114 ms, flip angle = 90°, matrix: 512 * 320, slice thickness: 2.5 mm, slice gap: 0.2 mm). The SBA cohort’s structural, T2-weighted MRI was performed with a fast recovery fast spin echo sequence (FRFSE, voxel dimensions = 0.7 × 0.7 × 2.7 mm^3^, TR = 5900 ms, TE = 97 ms, flip angle: 90°, matrix: 512 * 320, slice thickness: 2.5 mm, slice gap: 0.2 mm) in axial, sagittal and coronal planes. Also, for the EPO cohort, T2-weighted images were acquired with a FRFSE sequence (voxel dimensions = 0.7 × 0.7 × 2.8 mm^3^, TR = 6600 ms, TE = 126 ms, flip angle: 90°, matrix: 512 * 320, slice thickness: 2.5 mm, slice gap: 0.3 mm).

The HB and SB studies both acquired axial diffusion tensor imaging (DTI) data using a pulsed gradient spin-echo echo-planar imaging sequence (TR = 3950 ms, TE = 90.5 ms (for HB study) and TE = 90 ms (for SB study), field of view=18 cm, matrix=128×128, slice thickness=3 mm) with 35 diffusion encoding gradient directions at a b-value of 700 s/mm^2^ and four b=0 images. The EPO study used a pulsed gradient spin-echo echo-planar imaging sequence, but the following acquisition parameter differed: TR = 9000 ms, TE = 77 ms, 21 diffusion encoding gradient directions at a b-value of 1000 s/mm^2^ and four b=0 images.

### 2.3 MRI preprocessing

We used the same image processing pipeline for all etiologic groups with the possibility of adaptations according to the specific needs of the given cohort, such as the use of specific MRI template images. The main purpose of the MRI processing was to enable anatomically constrained probabilistic tractography (ACT) for whole-brain connectomic mapping to be carried out for the neonatal subjects using commonly reported procedures (Al Harrach et al., 2021; Batalle et al., 2017; Sa de Almeida et al., 2021), as well as to accommodate steps that were necessary to mitigate the specific challenges emerging from our newborn MRI acquisition. The pipeline was written in Python language (version 3.8) and wraps several algorithms that are commonly used to process MRI data, as described in the following paragraphs and in Figure 1 (https://github.com/annspe/connectome_pipeline).

The axial, sagittal and coronal structural T2-weighted images were super-resolution reconstructed into a single dataset using the SVRTK algorithm (Kuklisova-Murgasova et al., 2012) which led to a resampled in-plane resolution of 0.35 mm.

DTI data first underwent denoising using the deep learning algorithm patch2self from the DIPY package in Python (Fadnavis et al., 2020). This autoencoder is trained to learn a mapping between noisy diffusion imaging data and the same dataset with artificially added random noise. In this manner, the underlying structure of the data which is less affected by the noise is learned. This mapping can then be applied to the raw diffusion data. Next, Gibbs ringing artefact suppression (Kellner et al., 2016) implemented in MRtrix3Tissue (https://3Tissue.github.io), a fork of MRtrix3 (Tournier et al., 2019) was applied to remove high-frequency oscillations. Subsequently, motion and image distortion correction from the FSL software version 6.00 (S. M. Smith et al., 2004) was conducted using slice-to-volume eddy correction with CUDA 8.0. Lastly, B1 bias field correction using Advanced Normalization Tools (ANTs) N4 (Tustison et al., 2010) was applied with MRtrix3Tissue.

### 2.4 Network construction

The super-resolution reconstructed T2 images underwent ANTs N4 bias field correction (Tustison et al., 2010), followed by segmentation into seven tissue types (cortical grey matter, deep grey matter, white matter, external cerebrospinal fluid (CSF), ventricles, brainstem, cerebellum) using an in-house algorithm based on a U-Net architecture (Payette et al., 2021). This U-Net was trained on images including normal appearing brains as well as spina bifida. This step was necessary to ensure that we have representations of 5 tissue types for the anatomically constrained tractography using the same algorithm across all our etiologic groups.

The Edinburgh Neonatal Atlas (ENA33), which is a backpropagation of the Desikan-Killiany adult brain atlas (Desikan et al., 2006) to neonatal brains, was used to parcellate the brain (Blesa et al., 2016). Anatomical correspondence between the DTI and the atlas space was achieved by using the super-resolution reconstructed T2-weighted image of the subject as an intermediate image. The corresponding T2-weighted template image of the Atlas was registered to subject’s T2 image using the diffeomorphic symmetric image normalization method (SyN) in ANTs (Avants et al., 2014). The secondary purpose of using this non-linear registration step was to partially mitigate the geometric distortions occurring in the EPI data, since we did not have the possibility to carry out *topup* distortion correction for our clinically acquired data. For the MRI of spina bifida newborns, we used an in-house developed neonatal spina bifida atlas (Speckert et al., 2024).

Next, both the tissue segmentations as well as the Atlas based parcellations were next registered from the subject’s T2 space to the DTI space (average B = 0 image as target) using TRSAA + SynOnly (Avants et al., 2014) methods.

As the cerebellum was at least partially missing in 40 out of 187 DTI scans due to the incomplete coverage of the whole brain during the MRI acquisition, the cerebellar atlas regions had to be excluded for all subjects from the analysis. Further, the lateral ventricles and corpus callosum were excluded from the atlas, as their inclusion would not contribute meaningfully as nodes in the structural brain connectome. Finally, this led to an amended atlas of 94 regions (see supplementary Table 1).

Tissue response function estimations for white matter and CSF (Dhollander et al., 2019) were averaged across random 15 subjects per study population using MRtrix3Tissue. Based on that the orientation distribution functions (ODFs) for tissue and free water were estimated using the multi-shell multi-tissue constrained spherical deconvolution algorithm in MRtrix3Tissue, treating the b-value 0 as a shell. Next, the ODFs were normalized for each subject to be able to obtain quantitative measures of density (Dhollander et al., 2021; Tournier et al., 2019). Further, anatomically constrained probabilistic tractography (R. E. Smith et al., 2012; Tournier et al., 2010) with dynamic seeding (R. E. Smith et al., 2015) was used to generate 10 million streamlines. To ensure that the streamlines’ densities within the white matter closely approximate the fiber densities estimated through the spherical deconvolution diffusion model, SIFT2 (R. E. Smith et al., 2015; Tournier et al., 2019) was used. Further, the fiber density SIFT2 proportionality coefficient (µ) was calculated for each subject to achieve inter-subject connection density normalization. Lastly, structural connectomes were calculated for each subject with radial search radius of 4 mm with regions from the amended ENA33 atlas and the sum of weighted streamlines (SIFT2* µ) connecting each region as edges, resulting in a 94x94 structural connectivity matrix.

**Figure 1.**
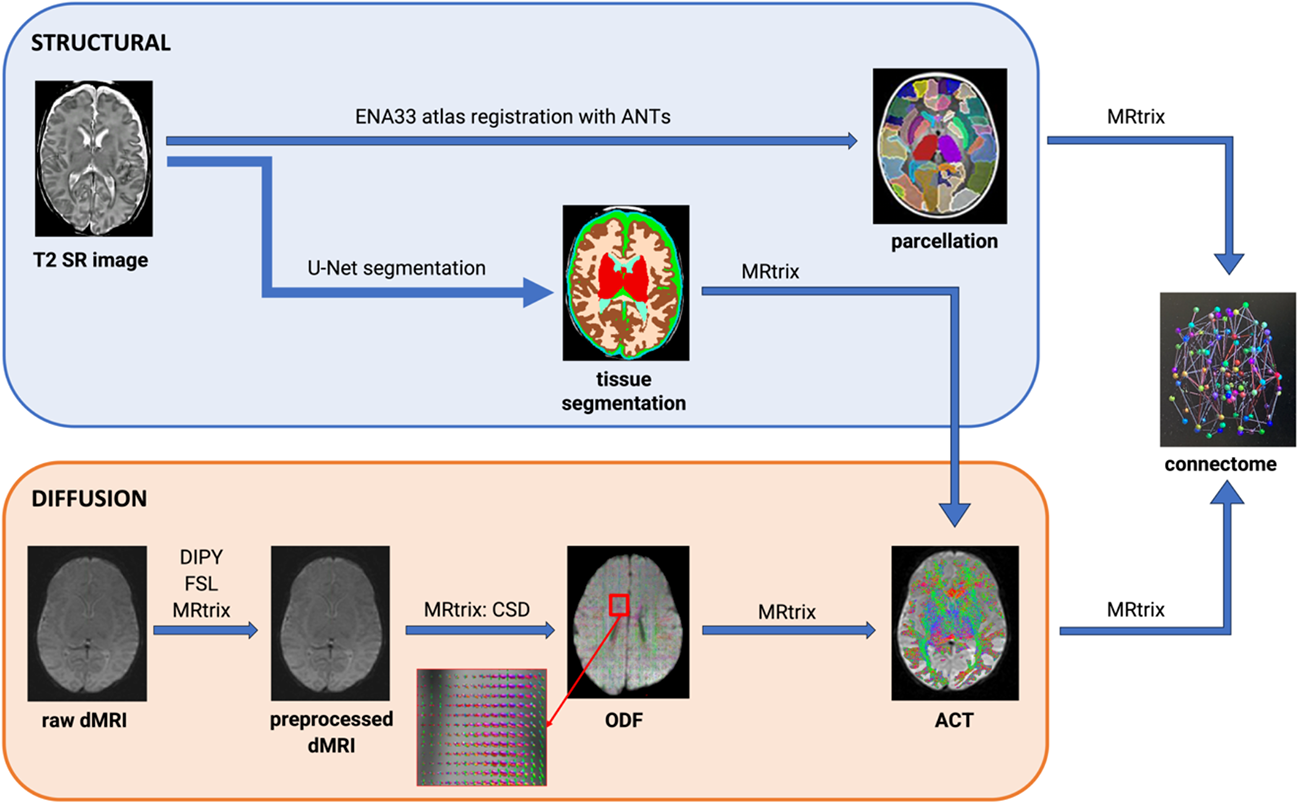
Connectome pipeline. *Note*. Overview of the image processing steps in the connectome pipeline which takes T2-weighted super-resolution and dMRI images as input and outputs a structural connectome. ANTs, DIPY, FSL and MRtrix are neuroimaging software. *dMRI* = diffusion MRI. *T2 SR* = T2-weighted super-resolution. *ENA33* = Edinburgh neonatal atlas. *CSD* = constrained spherical deconvolution. *ODF* = orientation distribution functions. *ACT* = anatomically constrained tractography.

### 2.5 Network measures and normalization using null-network models

Graph theoretical analyses were carried out to answer the hypothesis and assess further topological properties of individual weighted, undirected networks. Functions from the Brain Connectivity Toolbox (BCT) (Rubinov & Sporns, 2010) for MATLAB (version R2017b) were used. Global features of brain networks were characterized by assessing integration, segregation, and rich-club coefficients. Network integration was measured by the averaged global efficiency across multiple cost density thresholds. Global efficiency was calculated by taking the inverse of the average shortest path length between nodes, which quantifies the relative parallel information exchange among distributed regions within the network (Ní Bhroin et al., 2020; Rubinov & Sporns, 2010). Network segregation was assessed using the modularity coefficient, which describes the degree to which the “network may be subdivided into delineated and nonoverlapping groups” (Rubinov & Sporns, 2010, p. 1061).

The rich-club coefficient refers to the observation that hubs (nodes with many connections) in a network tend to be more densely interconnected with each other compared to nodes with lower degrees. Networks exhibiting rich clubs often demonstrate an efficient and higher-order organizational structure (Dennis et al., 2013).

As an additional global feature, we explored small-worldness, because it assesses the balance between segregation and integration within a network. It is defined as the ratio between the normalized clustering coefficient, representing segregation, and the normalized characteristic path length, representing integration. Small-world networks are positioned between random networks and so-called lattice networks, which show a high degree of clustering and a long characteristic path length (Miraglia et al., 2015). However, the definition does not provide an unambiguous quantitative method for implementation. We adopted the methodology introduced by (Muldoon et al., 2016) because of its suitability for weighted networks and its agnosticism to network density. The latter characteristic proves particularly crucial in our comparative analysis of networks with varying densities. For this analysis, MATLAB version R2023b was used due to the dependency of this function on the Bioinformatics Toolbox.

Furthermore, to further explore the topological properties of the networks, we additionally calculated the following local metrics: clustering coefficient, local efficiency, modularity coefficient, nodal strength, and betweenness-centrality. Local efficiency, which is another measure for segregation, evaluates the effectiveness of information exchange among adjacent nodes in the network (Ní Bhroin et al., 2020). The degree of an individual node is the number of edges connected to that node (Rubinov & Sporns, 2010). In weighted networks, nodal strengths, calculated as the sum of the connectivity weights associated with each node, offer additional insights into the network’s organization (Fornito et al., 2016). Betweenness-centrality, defined as the fraction of all shortest paths in the network passing through a given node, identifies bridging nodes connecting disparate parts of the network (Rubinov & Sporns, 2010).

To be able to further interpret the local characteristics of the network, we divided the structural connectivity networks into two distinct groups of nodes, core, and periphery nodes. This partitioning was achieved using the Kernighan-Lin algorithm for graph partitioning (Borgatti & Everett, 1999; Newman, 2006) available in the BCT toolbox (Rubinov & Sporns, 2010). In the resulting partitioning, core nodes exhibit strong connections with other core and periphery nodes, while periphery nodes display weaker interconnections with each other. A common core/periphery structure was defined such that the structure partitioned 90% of the subjects (see supplementary Table 1). An additional coreness statistic was computed, to estimate the degree of separation between core and peripheral nodes (Borgatti & Everett, 1999). Next, the local efficiency of core and peripheral connections was analyzed.

To enable meaningful comparisons of graph measures across the different etiologies, we employed a normalization method using null-network models (Váša & Mišić, 2022). These models provide a robust framework for benchmarking and comparing graphs by generating random equivalents for each connectome. Specifically, we utilized a widely recognized strategy known as the random wiring null-model, which ensures the preservation of key structural properties, such as the degree distribution, node numbers, and the number of edges (Chen et al., 2018). The BCT toolbox was used to generate 1000 random null-networks for each subject. For each subject’s global and local metrics, a Z-score was computed by normalizing the given metrics relative to the distribution of the metric values of the random null-networks corresponding to the same subject.

### 2.6 Low-dimensional representation of the connectomes

The visualization of the connectomes using the morphospace concept was carried out by representing them in a coordinate system established by the averaged global efficiency (measure of integration) and modularity coefficient (measure of segregation), based on the framework proposed by van den Heuvel & Sporns (2019). Next, to allow an embedding of the connectomes into a low-dimensional space, a local feature vector was created that combines local metrics of integration and segregation, serving as a comprehensive descriptor of each node’s contribution to the network. We used 1000 null-models (as described in 2.5) per case to calculate z-normalized local metrics. For each node, five metrics were calculated at 9 cost density thresholds from 0.1 to 0.9: clustering coefficient, local efficiency, modularity, nodal strength, and betweenness-centrality, leading to 45 features per sample. In instances where z-normalization resulted in ‘Inf’ or ‘-Inf’ values due to very low standard deviations, these occurrences were mitigated by substituting the affected entries with zero. This adjustment ensured the reliability of the z-normalized feature values for subsequent analysis.

To gain a visual understanding of how these local features differentiate between groups and relate to ND outcome, we employed a dimensionality reduction technique, specifically t-distributed stochastic neighbor embedding (t-SNE) (Van Der Maaten & Hinton, 2008). To achieve improved reproducibility, we incorporated the first three principal components as the initial position for t-SNE embedding. Combining principal component analysis (PCA) and t-SNE in this way is a common strategy to balance the non-linearity capturing abilities of t-SNE with a more deterministic starting point for improved consistency across runs (Becht et al., 2019; Kobak & Berens, 2019). The objective of graph embedding is to represent graphs in a vector space capturing differences and similarities among the original graphs (Bahonar et al., 2021). Thus, this approach facilitates the exploration of patterns in lower-dimensional space, offering an intuitive representation of the connectome’s local subtleties. By examining these local feature embeddings, we aimed to identify patterns that may underlie group distinctions and correlate with ND outcome.

### 2.7 Network-based statistics

To identify potential subnetworks of altered connectivity, we utilized network-based statistics (NBS) toolbox (Zalesky et al., 2010) for MATLAB (version R2017b). NBS detects subnetworks which differ between groups by permutation testing while considering multiple comparisons (Zalesky et al., 2010). When comparing the total number of connections between groups, we used a general linear model (GLM) with 10’000 permutations and multiple comparison correction with p = 0.05. The adjustment for multiple comparisons is achieved by cluster-based thresholding, wherein interconnected components within a network are regarded as a cluster. Our approach involved setting a primary test-statistics threshold at t = 3.1, which identifies a collection of supra-threshold connections where connections with a test statistic surpassing this threshold are deemed significant. Given the considerable impact of the primary test-statistics threshold on NBS outcomes, we systematically explored a spectrum of values (t = 2.5–3.5). The covariates for this analysis included gestational age (GA) at birth and postmenstrual age (PMA) at scan.

As NBS analysis relies on the values within connectivity matrices, direct group comparisons using this method were impractical. Consequently, NBS analysis was executed separately for each etiological group, with the dichotomized ND outcome serving as the grouping factor. The dichotomization employed a cut-off of one standard deviation below the mean, distinguishing between low and high outcomes.

### 2.8 Neurodevelopmental assessment

Children with CHD from the HB study had an ND assessment at 1 year of age and children with SBA at two years of age using the Bayley Scales of Infant and Toddler Development 3^rd^ edition (BSID-III) (Bayley, 2005), of which the Cognitive Composite Scores (BCCS) was used for this analysis, using the US normative reference data. Preterm-born children from the EPO study underwent testing at two years of age using the Bayley Scales of Infant Development 2^nd^ edition (BSID-II) Scores. For the present study, we used the mental development index (MDI), which was converted to the BCCS according to Jary et al. (2013). The test was administered by developmental pediatricians. BCCS was dichotomized at -1SD below the population mean, which lies at 92 BCCS.

### 2.9 Statistical analysis - general considerations

For group comparisons of sample characteristics, t-tests were utilized for normally distributed continuous data, Kruskal-Wallis test were utilized for non-normally distributed continuous data, and the Chi-squared test for categorical data.

Group differences in graph features, including integration, segregation, and rich club coefficient, were assessed through one-way analysis of covariance (ANCOVA). Covariates included gestational age (GA) at birth and postmenstrual age (PMA) at scan, with group (CHD, SBA, PB, healthy control) as the between-subject factor. Subsequently, a post-hoc analysis was conducted using Bonferroni correction to identify specific group differences.

To investigate the association between graph features and ND outcome, linear and logistic regression models were employed. Group (CHD, SBA, PB, healthy control) was treated as the predictor, and the BCCS were the dependent variable. For logistic regression, cognitive outcomes were dichotomized, again based on a cut-off of one standard deviation below the mean representing low outcome as opposed to high outcome. To investigate the core-periphery structure between the high and low ND outcome, Mann-Whitney U tests were employed. The same covariates as in the ANCOVA (GA and PMA) were included in these models.

Statistical analyses were carried out in R version 4.2.3. Throughout all the analysis, the significance level (alpha) was set at 0.05 and the assumptions for regression models and ANCOVA analysis were checked. Effect sizes were computed using Cohen’s d for t-tests and partial η^2^ for ANCOVAs.

## 3 Results

### 3.1 Descriptive statistics

The analysis irrespective of developmental outcome included 187 neonates, which comprised 42 healthy controls, 51 CHD, 51 PB and 43 SBA. Significant differences in gestational age at birth (GA) (p < 0.0001, Kruskal-Wallis test H(3) = 155.23) and postmenstrual age at scan (PMA) (p < 0.0001, Kruskal-Wallis test H(3) = 97.50) were revealed across the four groups. Post-hoc Dunn tests with Bonferroni correction for GA at birth indicated that only the only the CHD and Control groups did not significantly differ (p = 1), while significant differences were observed between the remaining pairs of groups (Control vs. PB p < 0.0001, Control vs. SBA p < 0.0001, CHD vs. PB p < 0.0001, CHD vs. SBA p < 0.0001, PB vs. SBA p = 0.0001). Regarding PMA at scan, the post-hoc Dunn test revealed that only the CHD and PB groups did not significantly differ (p = 0.36), while significant differences were observed between the remaining pairs of groups (Control vs. CHD p < 0.0001, Control vs. PB p < 0.0001, Control vs. SBA p < 0.0001, CHD vs. SBA p < 0.0001, PB vs. SBA p < 0.0001). Furthermore, the chi-square test indicated a significant difference in sex distribution between the groups (Chi-square = 12.775, df = 3, p = 0.005). Pairwise proportionality tests with Bonferroni correction revealed a significant difference in sex distribution only between the CHD and SBA groups (p = 0.014). Also, the BCCS differed across the four groups (p < 0.0001, ANOVA F(3,174) = 15.63). Post-hoc test with pairwise comparisons between groups with the Bonferroni correction revealed that the PB group did not significantly differ from the CHD (p = 1) and SBA (p = 0.27) groups, while the remaining pairs of groups differed (Control vs. CHD p = 0.0005, Control vs. PB p < 0.0001, Control vs. SBA p < 0.0001, CHD vs. SBA p = 0.02). Additional details about the sample characteristics can be found in Table 1.

**Table 1.** Sample characteristics of the study populations.

| Sample characteristic | Control<br>n = 42 | CHD<br>n = 51 | PB<br>n = 51 | SBA<br>n = 43 | p-value |
| --- | --- | --- | --- | --- | --- |
| GA at birth<br>(Mdn[IQR]) | 39.7<br>(37.5-41.9) | 39.4<br>(37.6-41.5) | 29.7<br>(27.6-31.8) | 36.0<br>(32.9-39.1) | $p < 0.0001^a$ |
| PMA at scan<br>(Mdn[IQR]) | 42.4<br>(40-44.8) | 40.3<br>(38.2-42.5) | 41.1<br>(37.8-44.3) | 38.0<br>(36.7-39.3) | $p < 0.0001^a$ |
| **Male sex (N[%])** | 19 (45.32%) | 36 (70.6%) | 31 (60.8%) | 16 (37.2%) | $p = 0.005^b$ |
| **BCCS (Mean[SD])** | 116.8 (11.5) | 106.1 (15.6) | 103.7 (8.5) | 98.6 (11.6) | $p < 0.0001^c$ |

*Note.* N = 187. Mdn = Median. IQR = Interquartile range. SD = Standard deviation. <sup>a</sup>Kruskal-Wallis test, <sup>b</sup>Chi-square test, <sup>c</sup>ANOVA test. *CHD* = congenital heart disease. *PB* = premature birth. *SBA* = spina bifida aperta.

### 3.2 Global network measures

One-way ANCOVA of z-normalized global network features revealed significant group differences in global efficiency measuring integration (F(3,182) = 7.66; p < 0.0001, η^2^ = 0.11), modularity measuring segregation (F(3,182) = 16.97; p < 0.0001, η^2^ = 0.22), and in mean rich-club coefficient (F(3,182) = 3.50; p = 0.017, η^2^ = 0.06) but not in small-worldness (F(3,182) = 2.58; p = 0.06, η^2^ = 0.04). In supplementary Table 2, one can find an overview of ANCOVA results including different covariates. Post-hoc analysis with Bonferroni correction showed that the z-normalized mean global efficiency score was significantly greater in premature babies (-6.29 +/-0.76) compared to healthy controls (-10.4 +/-0.46) and CHDs (-10.5 +/-0.47). Additionally, the mean global efficiency score was significantly higher in SBA (-7.07 +/-0.37) compared to healthy controls, and CHDs (Figure 2, top left). Further, the z-normalized mean modularity score was significantly greater in CHDs (16.7 +/-0.48) compared to SBA (12.7 +/-0.37) and PB (13.3 +/-0.78). SBA newborns showed lower mean modularity than healthy controls (15.6 +/-0.46) (Figure 2, top right). Further, the z-normalized mean rich club coefficient was found to be significantly greater in SBA (0.77 +/-0.18) compared to healthy controls (-0.18 +/-0.22) as can be seen in Figure 2 on the bottom left. Lastly, the z-normalized mean small-worldness was significantly smaller in CHDs (0.66 +/-0.04) compared to healthy controls (0.80 +/-0.04) (Figure 2, bottom right).

**Figure 2.**
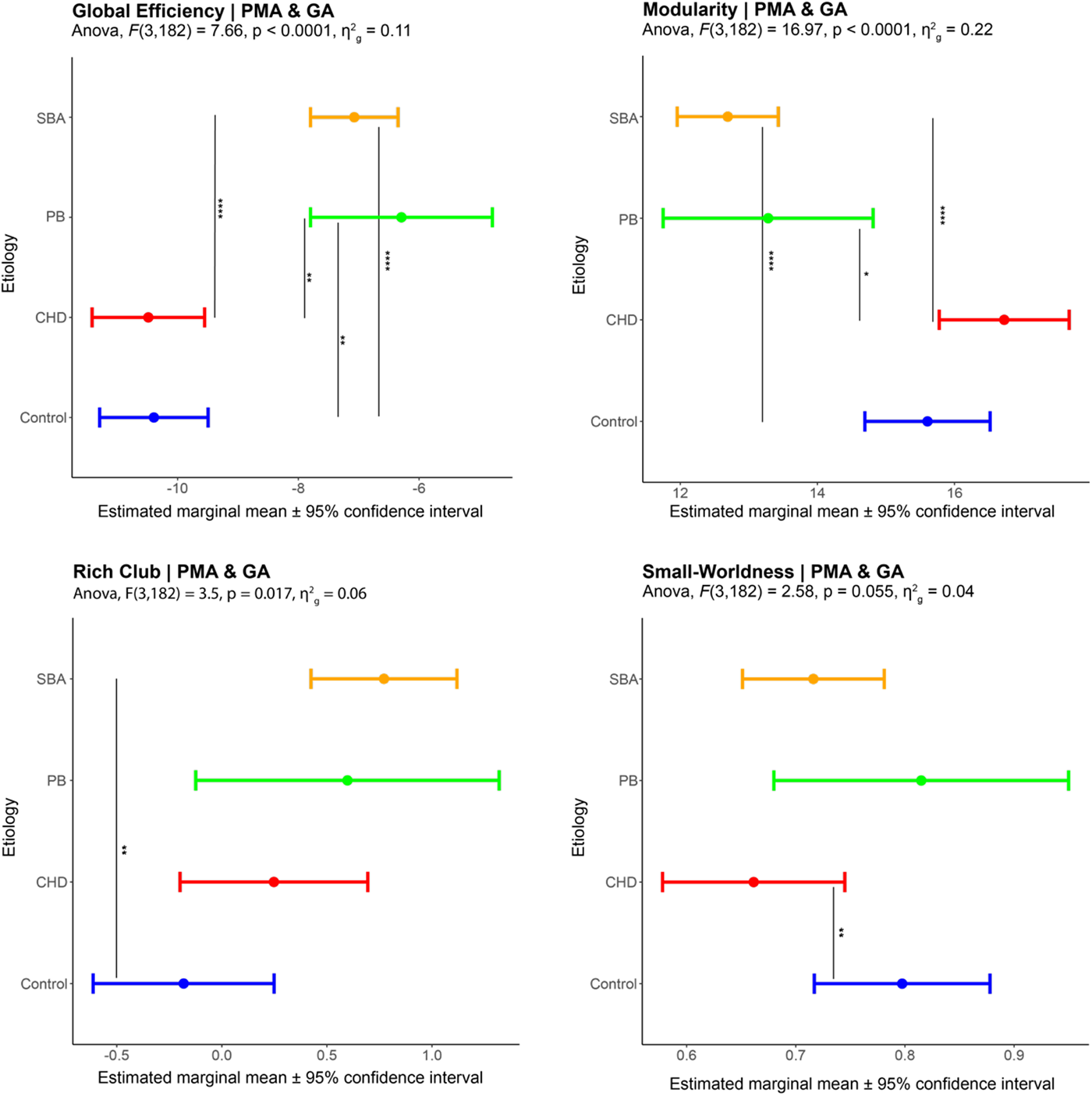
Adjusted means after post-hoc analysis of z-normalized global efficiency, z-normalized modularity, z-normalized rich club coefficients and small-worldness in different groups. *Note.* Pairwise comparison between groups using the emmeans test with Bonferroni correction showing the adjusted, estimated means. The colours depict the four different groups and the * flag the level of significance of the adjusted p-values . *SBA =* spina bifida aperta; *PB =* premature birth; *CHD =* congenital heart disease.

### 3.3 Low-dimensional representation of the connectomes

We employed two different methods to investigate the low-dimensional representations of the connectomes. Once within the integration-segregation morphospace based on the global features, and once with a t-SNE embedding of local features. As can be seen in Figure 3 on the left side, differences in the connectomic structure across the four groups are marginally visible. While the PB cohort and the controls highly overlap, the SBA cohort and the CHDs show more diverging profiles. Next, the t-SNE embedding (Figure 3, right side) shows three discernible patterns among the four groups: the SBA, the PB, and the HB cohort.

**Figure 3.**
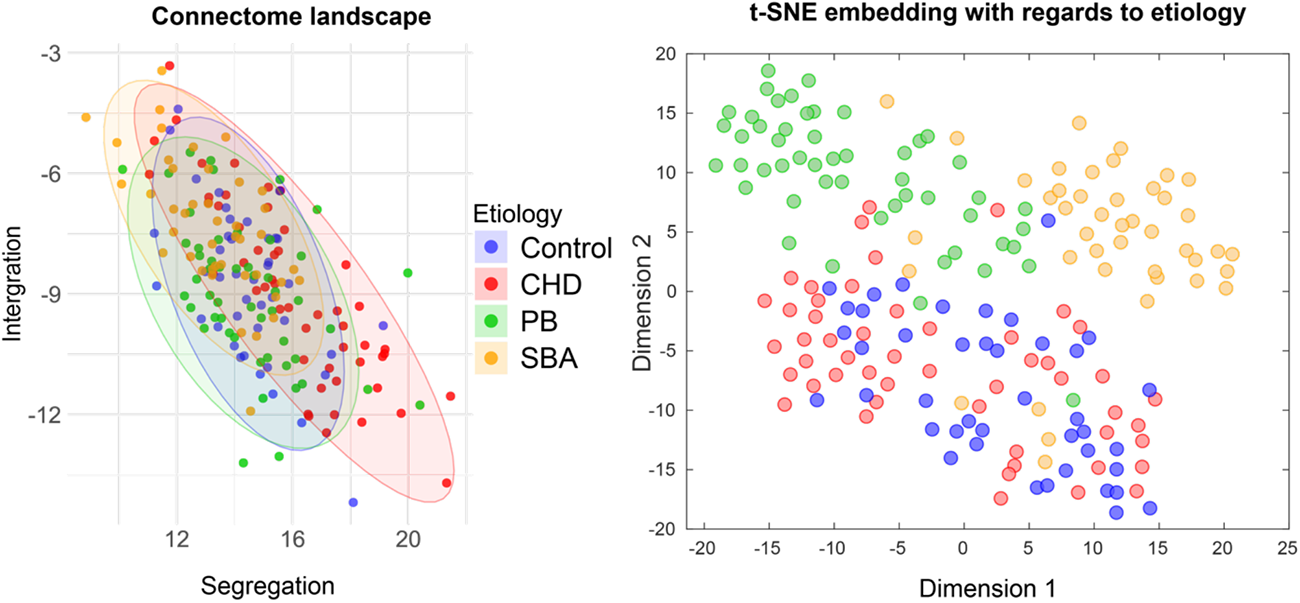
Two low-dimensional representations of the connectomes with respect to the four groups. *Note*. In the integration-segregation morphospace on the left side, the x- and y-axis show the z-normalized graph feature scores. In the t-SNE embedding on the right side, the initial positions were determined using the first 3 principal components for dimensionality reduction (Perplexity = 20; Standardize = True; Algorithm = exact) on the z-normalized local graph feature. *CHD =* congenital heart disease. *PB =* premature birth. *SBA =* spina bifida aperta.

### 3.4 Core-periphery structure

First, a common core/periphery structure (see supplementary Table 1 for a comprehensive list) was established for the entire study cohort, following the methodology outlined in section 2.5. Out of the 94 regions, 43 core nodes were identified which included the insula, precuneus, superior frontal cortex, and 32 periphery nodes. For each subject the coreness statistic was calculated. To investigate the core/periphery structure across etiologies, one-way ANCOVA of the z-normalized coreness revealed significant group differences (F(3,182) = 41.72, p < 0.0001, η^2^ = 0.41). Post-hoc analysis with Bonferroni adjustment showed that the SBA cohort (-1.89 +/-0.04) had significantly lower coreness than healthy controls (-1.40 +/-0.05), CHD (-1.32 +/-0.05), and PB (-1.57 +/-0.08).

Second, z-normalized local efficiency was calculated for the core and peripheral nodes separately to assess group differences. The mean local efficiency of the core (F(3,182) = 34.9, p < 0.0001, η^2^ = 0.37) and periphery (F(3,182) = 6.05, p < 0.001, η^2^ = 0.09) nodes significantly differed between the four groups. Also, after outlier removal (defined as 3 scaled median absolute deviations away from the median) the groups significantly differed (core nodes: F(3,181) = 38.46, p < 0.0001, η^2^ = 0.39; periphery nodes: F(3,177) = 4.75, p = 0.003, η^2^ = 0.08) as can be seen in Figure 4. Post-hoc analysis after outlier removal with Bonferroni adjustment for the core nodes showed that the SBA cohort (1.27 +/-0.05) had significantly lower mean local efficiency than the CHDs (1.77 +/-0.06), healthy controls (1.86 +/-0.06) and PB (1.59 +/-0.09). Similarly, for the periphery nodes, post-hoc analysis after outlier removal with Bonferroni correction showed that the SBA cohort (3.87 +/-0.27) has significantly higher mean local efficiency than the CHDs (2.71 +/-0.34) and healthy controls (2.51 +/-0.33).

**Figure 4.**
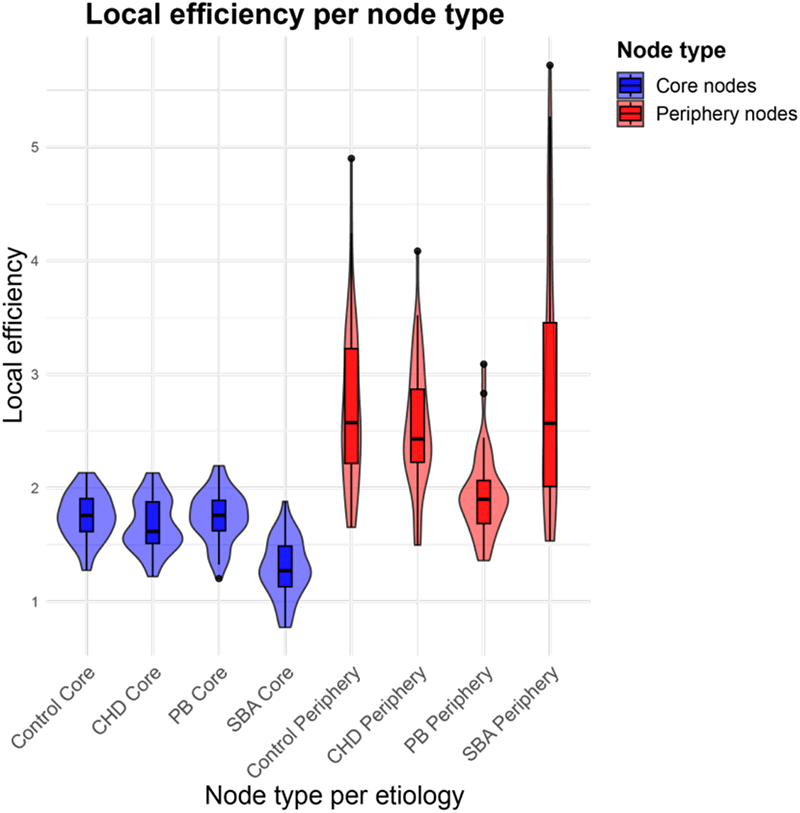
Average z-normalized local efficiency of core and periphery nodes across etiologies after outlier removal. *Note*. CHD = congenital heart disease. *PB =* premature birth. *SBA =* spina bifida aperta.

### 3.5 Association with neurodevelopmental outcome

The next step in our analysis aimed to assess the association of connectome parameters with ND outcome.

#### 3.5.1 Regression analysis

Linear regression analyses revealed a significant association between small-worldness and the BCCS (F(6, 171) = 10.02, adjusted R^2^ = 0.23, p < 0.0001, β_small-worldness_ = -12.05, p_small-worldness_ = 0.025). However, no statistically significant associations for neither integration (β = -0.14, p = 0.77), segregation (β = 0.05, p = 0.91) nor rich-club coefficient (β = 0.11, p = 0.917) was found. Subsequent linear regression analyses specific for each etiologic group revealed a significant association between small-worldness and cognitive outcome in the CHD cohort (F(3, 45) = 5.93, adjusted R^2^ = 0.24, p = 0.002, β_small-worldness_ = - 32.12, p_small-worldness_ = 0.005) as can be seen in Figure 5, while no such associations were observed for the PB and SBA cohorts etiology (see Table 2). Also, after BCCS outlier (n=2) removal the linear regression remained statistically significant (F(3, 43) = 6.95, adjusted R^2^ = 0.28, p = 0.0006, β_small-worldness_ = -28.76, p_small-worldness_ = 0.003) with a partial regression coefficient of r = -0.36 (p = 0.005).

**Table 2.**
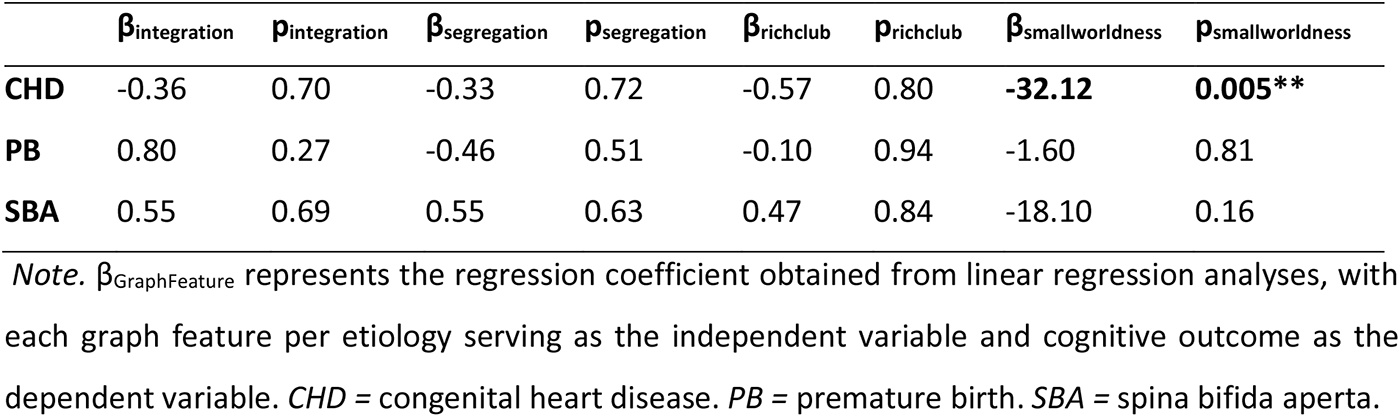
Linear regression results per etiology.

**Figure 5.**
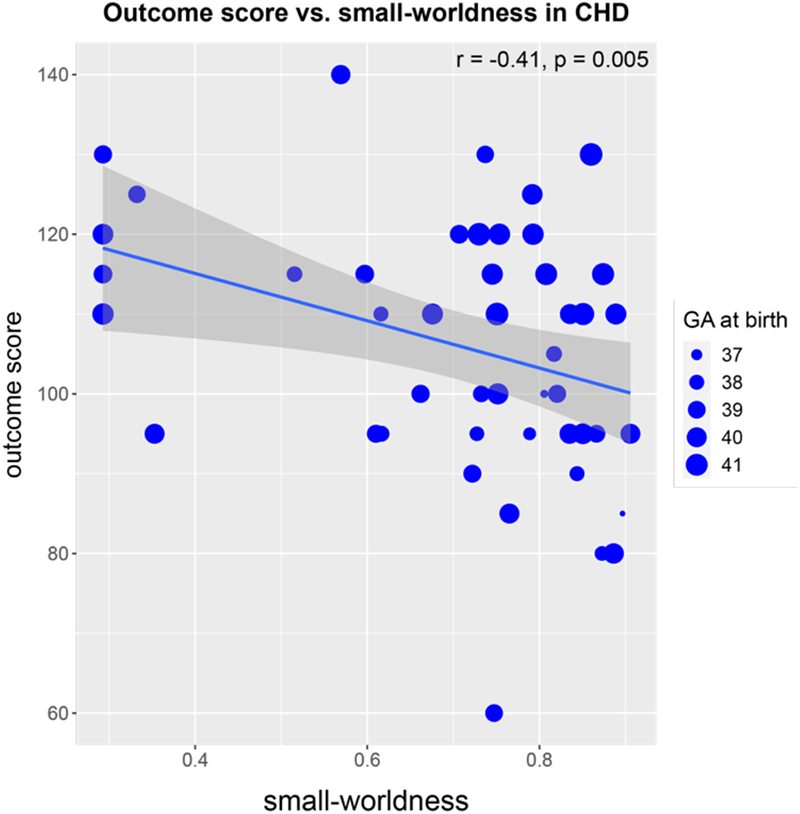
Scatterplot showing the negative relationship between small-worldness and neurodevelopmental outcome for the CHD cohort. *Note.* GA = gestational age. r = partial correlation coefficient.

In logistic regression analyses using dichotomized BCCS as the dependent variable, small-worldness persisted as the only predictor with a significant association across all subjects (β = 4.99, p = 0.03*). Integration (β = 0.18, p = 0.15), segregation (β = -0.17, p = 0.15) and rich-club coefficient (β = -0.14, p = 0.58) did not show significant associations. Further logistic regression analysis by etiology did not show any significant association between graph features and cognitive outcome in neither of the groups as can be seen in Table 3.

**Table 3.** Logistic regression results per etiology.

| | $\beta_{\text{integration}}$ | $p_{\text{integration}}$ | $\beta_{\text{segregation}}$ | $p_{\text{segregation}}$ | $\beta_{\text{richclub}}$ | $p_{\text{richclub}}$ | $\beta_{\text{smallworldness}}$ | $p_{\text{smallworldness}}$ |
| --- | --- | --- | --- | --- | --- | --- | --- | --- |
| **CHD** | 0.26 | 0.16 | -0.24 | 0.18 | -0.10 | 0.84 | 11.57 | 0.08 |
| **PB** | 0.11 | 0.71 | -0.06 | 0.84 | -0.14 | 0.81 | 2.67 | 0.43 |
| **SB** | -0.24 | 0.36 | -0.09 | 0.66 | -0.07 | 0.86 | 4.06 | 0.25 |

*Note.* $\beta_{\text{GraphFeature}}$ represents the regression coefficient obtained from logistic regression analyses, with each graph feature per etiology serving as the independent variable and cognitive outcome as the dependent variable. *CHD* = congenital heart disease. *PB* = premature birth. *SBA* = spina bifida aperta.

#### 3.5.2 Low-dimensional representation of the connectomes

We visualized the low-dimensional representation of the connectomes with regards to the dichotomised outcome scores. As can be seen in Figure 6, neither the the integration-segregation morphospace (left panel) based on the global features showed a discernable pattern regarding the dichotomized ND outcome, nor the t-SNE embedding of local features (right panel).

**Figure 6.**
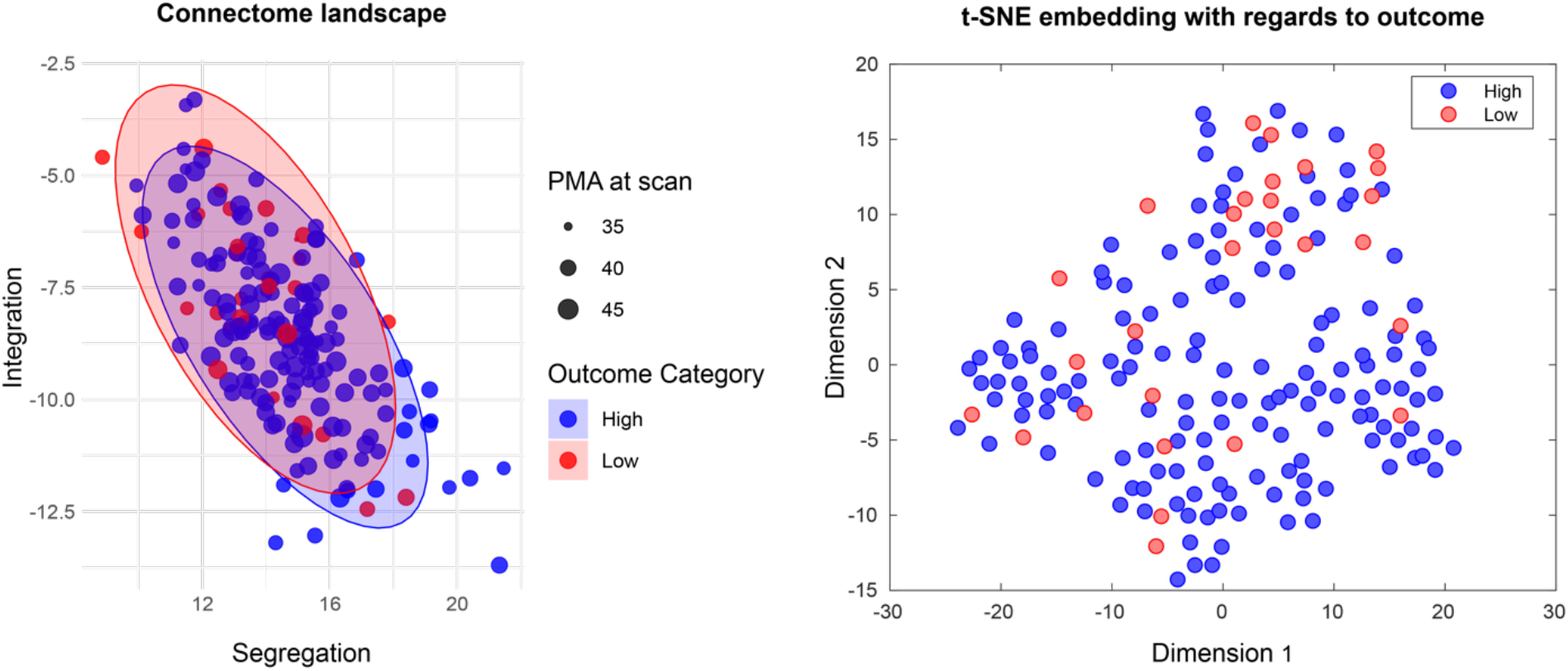
Two-dimensional connectomic landscape with respect to the neurodevelopmental outcome. *Note*. In the integration-segregation morphospace on the left side, the x- and y-axis show the z-normalized graph feature scores. In the t-SNE embedding on the right side, the initial positions were determined using the first 3 principal components for dimensionality reduction (Perplexity = 20; Standardize = True; Algorithm = exact) on the z-normalized local graph features). Low and high correspond to the dichotomized ND outcome scores from the BCCS.

#### 3.5.3 Core-periphery structure

The z-normalized local efficiency between high (median = 1.65, IQR = 0.38, n = 150) and low (median = 1.47, IQR = 0.38, n = 28) ND outcome differed significantly (W = 2798, p = 0.005, effect size r = 0.21) for the core nodes, but not for the periphery nodes (W = 2567, p = 0.06). This analysis was repeated separately for the etiologies, without significant results (CHD core nodes: W = 173, p = 0.47; CHD periphery nodes: W = 183, p = 0.32; PB core nodes: W = 135, p = 0.94; PB periphery nodes: W = 162, p = 0.39; SBA core nodes: W = 214, p = 0.79; SBA periphery nodes: W = 259, p = 0.15).

#### 3.5.4 Network-based statistics

Using NBS, no subnetworks could be identified which differed between the high and low ND outcomes within none of the four groups.

## 4 Discussion

To better understand the neural correlates of developmental delay in neonates at risk, our study examined the connectomic landscape with respect to the interplay between integration and segregation. Investigating a single institutional, cross-etiologic cohort, our study revealed connectomic differences among neonates with CHD, SBA, premature birth, and healthy controls. While observing shifts of these etiologies towards less optimal regions within the integration-segregation morphospace—driven by global graph features—we found no distinct, delineated disease-zones specific to either etiologies or ND outcomes. The application of t-SNE embedding of local graph features describing integration and segregation revealed a more discernible pattern of three distinct groups: PB, SBA, and a combined group of CHDs and healthy controls. No such pattern was evident regarding ND outcome, suggesting that local connectomic features describing integration and segregation in the brain can discern etiologic groups but not outcome groups. Despite the absence of clear disease zones in the morphospace, small-worldness emerged as a significant global graph feature. Its negative association with cognitive outcome, particularly within the CHD cohort, emphasizes the critical role of balancing segregation and integration in shaping neurodevelopmental trajectories.

### 4.1 Global network measures

We observed differences in global network measures across CHDs, SBA, PB, and healthy controls. Specifically, the SBA cohort exhibited higher z-normalized global efficiency compared to the CHDs and healthy controls (Figure 2, left), lower z-normalized modularity than CHDs and healthy controls (Figure 2, right), and higher mean z-normalized rich-club coefficient than healthy controls (Figure 2, bottom left). The CHD cohort displayed lower z-normalized global efficiency than PB (Figure 2, left), higher z-normalized modularity (Figure 2, right) than healthy controls and PB, and lower small-worldness than healthy controls (Figure 2, bottom left). Further, PB exhibited higher z-normalized global efficiency (Figure 2, left) than the healthy controls.

Our analysis used null-models for z-normalization, making direct comparisons with absolute, non-normalized features reported in the literature more challenging. The finding that pre-operative CHDs exhibit decreased segregation compared to healthy controls aligns with a report based on data taken from the same prospective cohort (Feldmann et al., 2020). Our finding of reduced global efficiency in CHDs aligns with Feldmann et al. (2020) findings of diminished integration in CHDs compared to healthy controls. Notably, while our study found significant differences between CHDs with prematurely born neonates, Feldmann et al. (2020) specifically compared CHDs to healthy controls. Despite the varied comparisons, both studies consistently indicate a decline in global efficiency among CHDs. Considering the widely proposed normative neonatal structural brain network development, where integration increases and segregation decreases over time (Jakab et al., 2014; Keunen, 2017; Song et al., 2017; Takahashi et al., 2012; Tymofiyeva et al., 2013), one could argue that CHDs experience delayed brain maturation compared to premature births. However, if we extend this perspective to the increased integration and decreased segregation observed in the SBA cohort compared to healthy controls, it suggests either an advanced stage of brain maturation of the SBA cohort or necessitates an alternative explanation. Similarly, the observed increased integration of the PB cohort compared to healthy controls and decreased segregation compared to CHD, raises the question of whether increased integration and decreased segregation indicate dysmaturation. Comparing our result with existing studies, we find similarities with by Zhao et al. (2019) and Kline et al. (2021) reporting a decrease in segregation compared to neonates with a higher gestational age at birth. Contrarily, finding increased segregation in PB is not supported by the literature. One possible reason for that is that prematurity encompasses a broad spectrum of gestational ages (< 37 weeks), and within this range, there can be considerable variation in neurodevelopmental trajectories. Factors such as degree of prematurity, severity of respiratory distress, and intrauterine growth restriction can influence brain development differently (Leijon, 2010). Therefore, the contradictory results observed in PB infants might stem from this heterogeneity within the prematurity population.

For small-worldness, our findings might contradict those of Schmithorst et al. (2018), who reported increased small-worldness in CHDs compared to healthy controls. It’s crucial to highlight the methodological differences underlying these contrasting results. Our approach, tailored for weighted networks independent of network density, is different from Schmithorst et al. (2018) method, which is not agnostic to network density and hence necessitated correction for the number of connections in each network. The importance of employing density-independent network analyses for effective network comparison is emphasized by the growing literature on neurodevelopment, disease differentiation, and aging (Muldoon et al., 2016). By mitigating the impact of network density on small-worldness computation, a more direct comparison of topological network structures between groups is enabled, minimizing the influence of density variations (Muldoon et al., 2016). A study by Hu et al. (2023), which employed the small-worldness computation suggested by Muldoon et al. (2016,) has found that the functional brain network in preterm born individuals showed considerable underdevelopment of small-world architecture compared to term-equivalent age or full-term born neonates. A study investigating the functional networks of young adults, deviations of small-world architectures have been observed in attention deficit hyperactivity (ADHD) and in schizophrenia (Liao et al., 2017). The observed decrease in small-worldness in CHDs aligns with the broader trend of small-world alterations in neurological and psychiatric diseases (Liao et al., 2017). Our findings contribute to this understanding by highlighting CHDs as having a decrease in small-worldness compared to healthy controls.

### 4.2 Low-dimensional representation of the connectomes

In the context of our cross-etiologic study, we are faced with the challenges of comparing network measures due to the absence of a ‘common space’ that could link connectome alterations across different disorders (van den Heuvel & Sporns, 2019). Therefore, we used null-models for network normalization (Váša & Mišić, 2022) as well as representations in a lower-dimensional space. In the two-dimensional integration-segregation morphospace, we revealed notable shifts across the four groups, yet the absence of distinct disease-specific zones suggests a more nuanced topological variation without clear-cut clusters (Figure 3, left). These observed shifts point towards a continuum of variations rather than discrete groupings. It becomes apparent that the connectomic differences may be more subtle and not easily captured by global graph features. In contrast, the application of t-SNE embedding to nodal graph features related to integration and segregation revealed a more discernible pattern (Figure 3, right). This implies that specific, especially regional aspects of integration and segregation play a pivotal role in distinguishing etiological groups.

### 4.3 Core-periphery structure

We identified a common core-periphery structure across all subjects (see supplementary Table 1). The identified core regions included insula, precuneus, and superior frontal cortex which is in line with previous descriptions in the neonatal literature (Ní Bhroin et al., 2020). We found that the SBA cohort exhibited a distinctive pattern – significantly decreased local efficiency in core nodes and increased local efficiency in periphery nodes, compared to CHDs and healthy controls, persisting also after outlier removal (Figure 4). Drawing on the work of Ball et al. (2014), core structures, established by the 30th gestational week, are prioritized during neurodevelopment over more peripheral connections (Jakab et al., 2019). This prioritization suggests that the observed network configuration in the SBA cohort might be indicative of a developmental delay, supported by the increased peripheral connections known to be associated with prematurity (Ball et al., 2014).

### 4.4 Association with neurodevelopmental outcome

Overall, linear regression analyses revealed a significant negative association between small-worldness and the BCCS in the combined group comprising all subjects. Specific within-etiology analyses revealed that only the CHD cohort demonstrated a significant negative association between small-worldness and cognitive outcomes (Figure 5). These results highlight the association between small-worldness in brain networks and neurodevelopmental outcomes, particularly within the CHD population. Our study therefore validates the hypothesis by Muldoon et al. (2016) that accounting for weak connections by using weighted measures instead of hard thresholding might reveal associations between small-worldness and cognition. Our study identified a significant association between small-worldness and cognition in these neonatal groups. These findings contribute to the growing body of evidence supporting the relevance of weak connections in predicting cognitive functioning. This aligns with a broader spectrum of research that has linked small-worldness to brain aging, neurodegeneration (Kukla et al., 2022), and Alzheimer’s disease (Zdanovskis et al., 2021).

In addition to finding significant associations of small-worldness with ND outcomes, we identified a small but significant effect of increased z-normalized local efficiency in the core nodes of subjects with high ND outcomes compared to those with low outcomes. As core nodes are prioritized during healthy neurodevelopment (Ball et al., 2014), increased local efficiency might be indicative of quicker brain maturation. Other analyses investigating the association to ND outcome, including identifying subnetworks using NBS and the exploration of lower-dimensional feature representations (Figure 6), failed to provide additional insights into ND outcomes as neither subnetworks could be identified, nor a clear separation was visually evident in Figure 6.

### 4.5 Limitations and future perspectives

We identified the following limitations in our study. Firstly, the inherent differences among the four groups, such as variations in gestational age at birth, postmenstrual age at scan, administration of sedation during scanning, and MRI parameters, present challenges in direct comparison, despite adjusting for these variables in our statistical models. To address this, we used a standardized preprocessing pipeline, z-normalization with null-models (Váša & Mišić, 2022) and we corrected for GA at birth and PMA at scan in our statistical models to enhance comparability of connectomic measures. Matching PMA at scan is common in case-control studies; however, the resulting variations in the time between birth and scan, particularly notable between healthy controls and preterm birth, pose a challenge. The rapid post-birth perfusion increase (Miranda et al., 2006) introduces potential bias in comparing different neonatal cohorts. In the future, additional work is necessary on larger cohorts, from which it is possible to sample a more balanced age distribution, or alternatively, to enroll cases with a more equal age distribution in follow-up prospective studies. Secondly, the characteristic brain morphology of the SBA group and to a less extent of the preterm group may introduce potential bias emerging from the different brain appearance and anatomy. This was partially mitigated by using an SBA-specific brain atlas in our analysis, aiming to ensure an anatomically more accurate localization of regions of interests despite the presence of ventriculomegaly and other abnormalities. Despite efforts to align SBA atlas nodes with the ENA33 atlas nodes, inherent morphological differences in SBA, such as increased ventricles, may limit the generalizability of our results in these two groups. Thirdly, the CHD cohort includes several subtypes of heart defects (Feldmann et al., 2020), introducing a potential influence of the cardiac physiology on brain development and structural brain networks. In future studies, larger CHD cohorts are needed to ensure each subtype is adequately represented, allowing for a separate examination of CHD subtypes. Fourthly, our study sample demonstrated high performance in ND outcomes, resulting in a high cut-off value for dichotomization, potentially limiting generalizability. Additionally, the age discrepancy in administering the BSID test between PB infants and other groups may introduce a bias. However, it is worth noting that the BSID test at 12 months has demonstrated predictive value for later cognitive development (Krogh & Væver, 2019).

Methodologically, the exclusion of the cerebellar nodes in the connectome construction might limit the comprehensive understanding of brain network connectivity. Future studies should be careful during image acquisition to ensure the cerebellum is adequately captured, thereby avoiding the need for exclusion, and enabling a more comprehensive examination of brain network connectivity. Moreover, the creation of the node feature vector for t-SNE embedding relies on our selection of features and thresholds, introducing a degree of subjectivity, however, future works should empirically determine the optimal set of features for such low-dimensional embedding, as well as carry out a comparison of different techniques. Future work should analyze longer-term cognitive outcomes, which might offer a more complete perspective on development (O’Shea et al., 2018).

### 4.6 Conclusion

Our study improves our understanding of the connectome’s duality in the context of newborns at risk for cognitive developmental delay. Our findings demonstrate that alterations in the interplay between modular specialization and efficient integration, especially assessed by small-worldness, are associated with both the neonatal connectomic topology and cognitive neurodevelopment.

## ACKNOWLEDGMENTS

The authors want to first thank all families who participated in this research. In addition, we thank our contributing study group without whom the spina bifida research would not be possible. From the University Children’s Hospital this includes Barbara Casanova, Ruth Etter, Domenic Grisch, Mirjam Harm, Maya Horst Luethy, Jenny Kienzler, Niklaus Krayenbuehl, Markus A. Landolt, Andreas Meyer-Heim, Theres Moehrlen, Svea Muehlberg, Evelyne Riesen, Brigitte Seliner, Mithula Shellvarajah, Alexandra Wattinger and Noemi Zweifel. From the University Hospital Zurich, the study group consists of Lukas Kandler, Nele Struebing, Max Antonio Thomasius and Ladina Vonzun. Infrastructure support for the spina bifida research was provided by the Clinical Trial Center, University Hospital of Zurich. The authors also want to thank the The Swiss EPO Neuroprotection Trial Group: (alphabetical order by study site): Aarau: Neonatal Unit, Department of Pediatrics, Kantonsspital Aarau (Georg Zeilinger, MD; Sylviane Pasquier, MD), Department of Neuropediatrics (A. Capone). Basel: Universitätskinderklinik UKBB (Christoph Bührer, MD; René Glanzmann, MD; Sven Schulzke, MD), Department of Neuropediatric and Developmental Medicine (P. Weber). Chur: Abteilung für Neonatologie, Kantons-und Regionalspital (Brigitte Scharrer, MD; Walter Bär, MD), Department of Neuropediatrics (E. Keller, C. Killer). Geneva: Neonatology and Pediatric Intensive Care, Department of Pediatrics, University Hospital HCUG (Riccardo Pfister, MD; Krämer Karin, MD), Division of Development and Growth (P.S. Hüppi, C. Borradori-Tolsa). Zurich (Hans Ulrich Bucher, MD; Jean-Claude Fauchère, MD; Brigitte Koller, MSc.; Sven Wellmann, MD; Cornelia Hagmann, MD, PhD), University Children’s Hospital Zurich, Child Development Centre (B. Latal, G. Natalucci). The EPO study was supported by a grant received from the Swiss National Science Foundation (3200B0-108176) and was registered at clinicaltrials.gov (identifier: NCT 00313946).

