## Supplementary Files for "Altered connectome topology in newborns at risk for cognitive developmental delay: a cross-etiologic study"

### Supplementary Material

#### Supplementary Figure 1: Flowchart of data inclusion of HB dataset

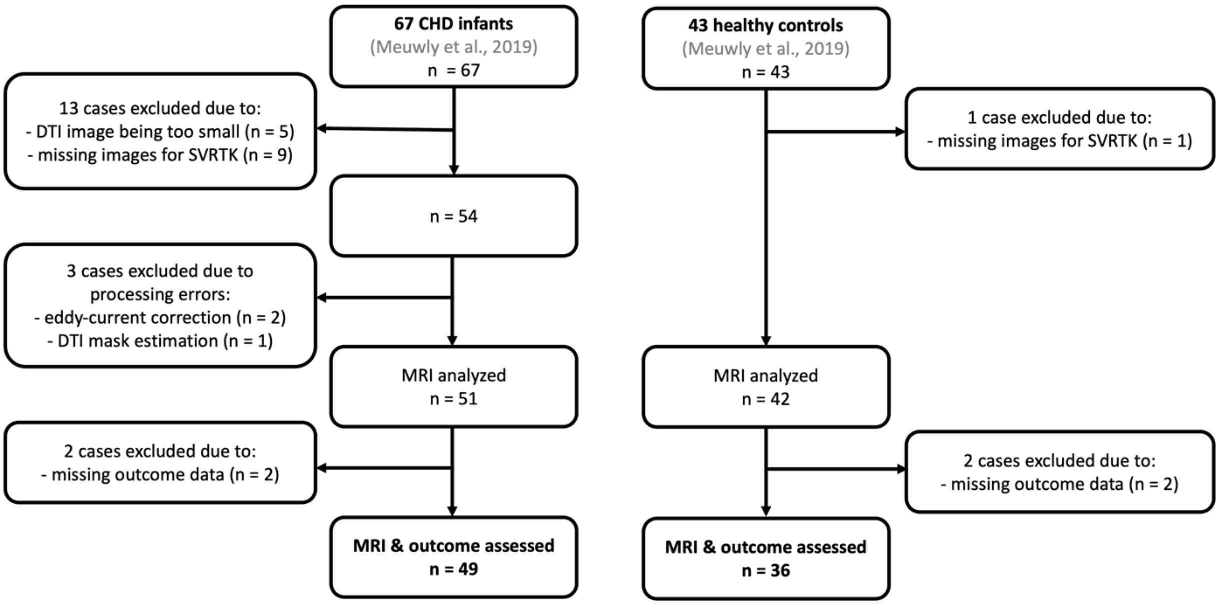


*Note.* The original inclusion and exclusion criteria can be found in Meuwly et al. (2019). CHD = congenital heart defect. DTI = diffusion tensor imaging. SVRTK = super-resolution reconstruction algorithm. MRI = magnet resonance image.

#### Supplementary Figure 2: Flowchart of data inclusion of SB and PB dataset

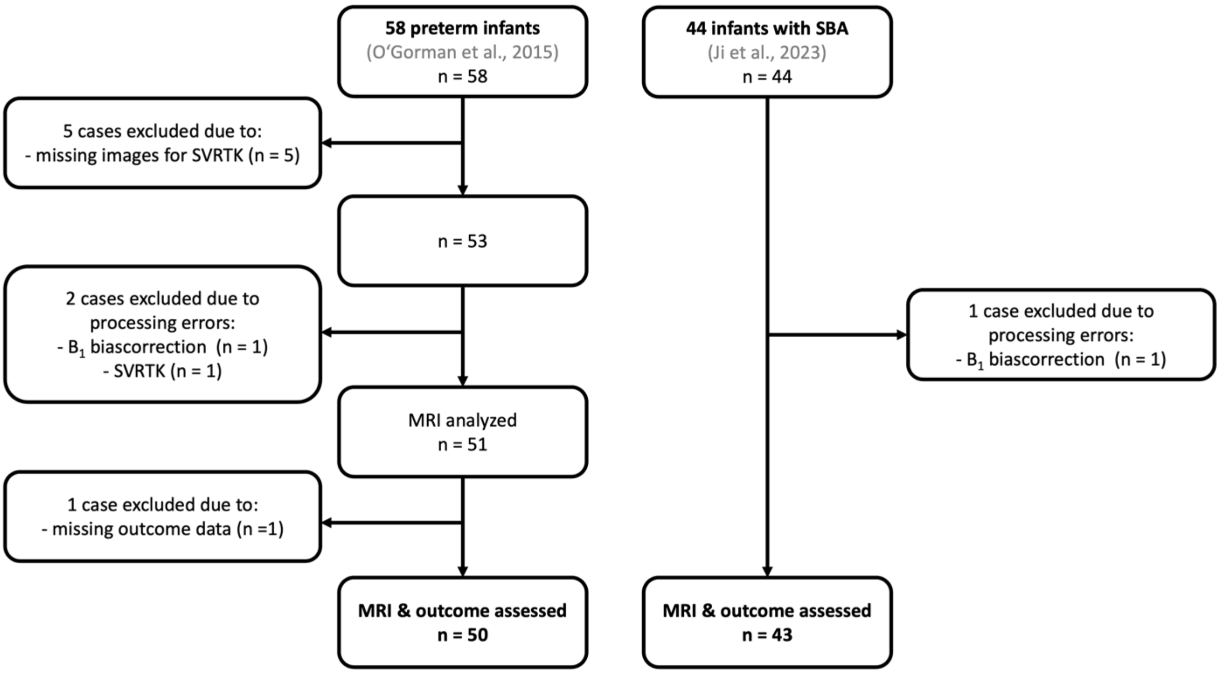


*Note.* The original inclusion and exclusion criteria can be found in O'Gorman et al. (2015) for the preterm infants and in Ji et al. (2023) for the SBA infants. SBA = spina bifida aperta. SVRTK = super-resolution reconstruction algorithm. MRI = magnet resonance image.

**Supplementary Table 1: Amended ENA33 atlas**

| LabelID | Anatomical Definition | Node Type | | LabelID | Anatomical Definition | Node Type |
|---|---|---|---|---|---|---|
| 1 | Precentral_L | Core | | 48 | Lingual_R | Core |
| 2 | Precentral_R | Core | | 49 | Occipital_Sup_L | NaN |
| 3 | Frontal_Sup_L | Core | | 50 | Occipital_Sup_R | Core |
| 4 | Frontal_Sup_R | Core | | 51 | Occipital_Mid_L | Core |
| 5 | Frontal_Sup_Orb_L | NaN | | 52 | Occipital_Mid_R | Core |
| 6 | Frontal_Sup_Orb_R | Periphery | | 53 | Occipital_Inf_L | NaN |
| 7 | Frontal_Mid_L | Core | | 54 | Occipital_Inf_R | Core |
| 8 | Frontal_Mid_R | Core | | 55 | Fusiform_L | Core |
| 9 | Frontal_Mid_Orb_L | Periphery | | 56 | Fusiform_R | Core |
| 10 | Frontal_Mid_Orb_R | Periphery | | 57 | Postcentral_L | Core |
| 11 | Frontal_Inf_Oper_L | NaN | | 58 | Postcentral_R | Core |
| 12 | Frontal_Inf_Oper_R | NaN | | 59 | Parietal_Sup_L | Core |
| 13 | Frontal_Inf_Tri_L | NaN | | 60 | Parietal_Sup_R | Core |
| 14 | Frontal_Inf_Tri_R | NaN | | 61 | Parietal_Inf_L | NaN |
| 15 | Frontal_Inf_Orb_L | Core | | 62 | Parietal_Inf_R | Core |
| 16 | Frontal_Inf_Orb_R | Core | | 63 | SupraMarginal_L | Core |
| 17 | Rolandic_Oper_L | Core | | 64 | SupraMarginal_R | NaN |
| 18 | Rolandic_Oper_R | Core | | 65 | Angular_L | Core |
| 19 | Supp_Motor_Area_L | Core | | 66 | Angular_R | NaN |
| 20 | Supp_Motor_Area_R | Core | | 67 | Precuneus_L | Core |
| 21 | Olfactory_L | Periphery | | 68 | Precuneus_R | Core |
| 22 | Olfactory_R | Periphery | | 69 | Paracentral_Lobule_L | Periphery |
| 23 | Frontal_Sup_Medial_L | Core | | 70 | Paracentral_Lobule_R | NaN |
| 24 | Frontal_Sup_Medial_R | Core | | 71 | Caudate_L | Periphery |
| 25 | Frontal_Med_Orb_L | Periphery | | 72 | Caudate_R | Periphery |
| 26 | Frontal_Med_Orb_R | Periphery | | 73 | Putamen_L | NaN |
| 27 | Rectus_L | Periphery | | 74 | Putamen_R | NaN |
| 28 | Rectus_R | Periphery | | 75 | Pallidum_L | Periphery |
| 29 | Insula_L | Core | | 76 | Pallidum_R | Periphery |
| 30 | Insula_R | Core | | 77 | Thalamus_L | NaN |
| 31 | Cingulum_Ant_L | NaN | | 78 | Thalamus_R | NaN |
| 32 | Cingulum_Ant_R | NaN | | 79 | Heschl_L | Periphery |
| 33 | Cingulum_Mid_L | Core | | 80 | Heschl_R | Periphery |
| 34 | Cingulum_Mid_R | Core | | 81 | Temporal_Sup_L | Core |
| 35 | Cingulum_Post_L | Periphery | | 82 | Temporal_Sup_R | Core |
| 36 | Cingulum_Post_R | Periphery | | 83 | Temporal_Pole_Sup_L | Periphery |
| 37 | Hippocampus_L | Periphery | | 84 | Temporal_Pole_Sup_R | Periphery |
| 38 | Hippocampus_R | Periphery | | 85 | Temporal_Mid_L | Core |
| 39 | ParaHippocampal_L | Periphery | | 86 | Temporal_Mid_R | Core |
| 40 | ParaHippocampal_R | Periphery | | 87 | Temporal_Pole_Mid_L | Periphery |
| 41 | Amygdala_L | Periphery | | 88 | Temporal_Pole_Mid_R | Periphery |
| 42 | Amygdala_R | Periphery | | 89 | Temporal_Inf_L | Core |
| 43 | Calcarine_L | Core | | 90 | Temporal_Inf_R | Core |
| 44 | Calcarine_R | Core | | 91 | Midbrain_L | Periphery |
| 45 | Cuneus_L | NaN | | 92 | Midbrain_R | Periphery |
| 46 | Cuneus_R | NaN | | 93 | Pons_L | Periphery |
| 47 | Lingual_L | Core | | 94 | Pons_R | Periphery |

**Supplementary Table 2: ANCOVA results including different covariates**

| | Global Efficiency | Global Efficiency | Global Efficiency | Global Efficiency |
|---|---|---|---|---|
| Etiology | F(3, 183) = 7.68; p < 0.0001*** | F(3, 182) = 7.66; p < 0.0001*** | F(3, 183) = 7.68; p < 0.0001*** | F(3, 182) = 7.66; p < 0.0001*** |
| PMA at scan | | x | | x |
| GA at birth | | | x | x |
| | Modularity | Modularity | Modularity | Modularity |
| Etiology | F(3, 183) = 16.18; p < 0.0001*** | F(3, 182) = 16.97; p < 0.0001*** | F(3, 183) = 16.18; p < 0.0001*** | F(3, 182) = 16.97; p < 0.0001*** |
| PMA at scan | | x | | x |
| GA at birth | | | x | x |
| | Rich Club | Rich Club | Rich Club | Rich Club |
| Etiology | F(3, 183) = 3.51; p = 0.016* | F(3, 182) = 3.50; p = 0.017* | F(3, 183) = 3.51; p = 0.016* | F(3, 182) = 3.50; p = 0.017* |
| PMA at scan | | x | | x |
| GA at birth | | | x | x |
| | Small-Worldness | Small-Worldness | Small-Worldness | Small-Worldness |
| Etiology | F(3, 183) = 2.59; p = 0.05 | F(3, 182) = 2.58; p = 0.06 | F(3, 183) = 2.59; p = 0.05 | F(3,182) = 2.58; p = 0.06 |
| PMA at scan | | x | | x |
| GA at birth | | | x | x |

*Note*. The homogeneity of regression slopes between etiology and the covariates was given, meaning that the interaction term in the linear models was not significant for either of the models. Therefore, only models without an interaction effect were analyzed and showed here.

**Supplementary Table 3: CHD types**

| | Subject with successful connectome creation n (%) | Subject with connectome and outcome n (%) |
|---|---|---|
| **Type of CHD** | | |
| dTGA | 36 (70.6) | 36 (73.5) |
| single ventricle | 6 (11.7) | 5 (10.2) |
| CoA | 4 (7.8) | 4 (8.2) |
| DORV | 1 (2.0) | 0 (0.0) |
| interrupted aortic arch | 1 (2.0) | 1 (2.0) |
| other | 3 (5.9) | 3 (6.1) |
| **total** | 51 | 49 |

Note. CHD = congenital heart disease; dTGA = d-transposition of great arteries; CoA = coarctation of the aorta; DORV = double outlet right ventricle. Single ventricle including Hypoplastic Left Heart Syndrome, Hypoplastic Left Heart Complex, Double Inlet Left Ventricle, Pulmonary Atresia with intact Ventricular Septum; Tricuspid Atresia, Left Atrial Isomerism. Other including Ventricular Septal Defect, Common Arterial Trunk, congenital corrected TGA (L-TGA).
